# Joint imputation and deconvolution of gene expression across spatial transcriptomics platforms

**DOI:** 10.1101/2025.02.17.638195

**Authors:** Hongyu Zheng, Hirak Sarkar, Benjamin J. Raphael

**Affiliations:** Department of Computer Science, Princeton University, Princeton, NJ, USA; Ludwig Cancer Institute, Princeton Branch, Princeton University, Princeton, NJ, USA

## Abstract

Spatially resolved transcriptomics (SRT) technologies measure gene expression across thousands of spatial locations within a tissue slice. Multiple SRT technologies are currently available and others are in active development with each technology having varying spatial resolution (subcellular, single-cell, or multicellular regions), gene coverage (targeted vs. whole-transcriptome), and sequencing depth per location. For example, the widely used 10x Genomics Visium platform measures whole transcriptomes from multiple-cell-sized spots, while the 10x Genomics Xenium platform measures a few hundred genes at subcellular resolution. A number of studies apply multiple SRT technologies to slices that originate from the same biological tissue. Integration of data from different SRT technologies can overcome limitations of the individual technologies enabling the imputation of expression from unmeasured genes in targeted technologies and/or the deconvolution of ad-mixed expression from technologies with lower spatial resolution. We introduce Spatial Integration for Imputation and Deconvolution (SIID), an algorithm to reconstruct a latent spatial gene expression matrix from a pair of observations from different SRT technologies. SIID leverages a spatial alignment and uses a joint non-negative factorization model to accurately impute missing gene expression and infer gene expression signatures of cell types from ad-mixed SRT data. In simulations involving paired SRT datasets from different technologies (e.g., Xenium and Visium), SIID shows superior performance in reconstructing spot-to-cell-type assignments, recovering cell-type-specific gene expression, and imputing missing data compared to contemporary tools. When applied to real-world 10x Xenium-Visium pairs from human breast and colon cancer tissues, SIID achieves highest performance in imputing holdout gene expression. A PyTorch implementation of SIID is available at https://github.com/raphael-group/siid.

## 1 Introduction

Spatially resolved transcriptomics (SRT) technologies have transformed the study of tissue biology by enabling the simultaneous measurement of gene expression at thousands of locations within a tissue section, with each expression measurement linked to its corresponding spatial coordinate. SRT technologies allow researchers to study complex spatial gene expression patterns and intricate cellular organization, providing a closer look at the tissue microenvironment and spatial context for a given disease [35, 33]. There are multiple SRT technologies currently in use with varying spatial resolution and breadth/depth of gene expression [31, 37]. For example, in situ capture based SRT such as Slide-seq [38, 36] and 10x Genomics Visium measure thousands of genes using barcoded beads (of radius 10–50*µm*) on a slide. Here, a single bead captures mRNA molecules from multiple spatially nearby cells and thus the gene expression measurement is for a mixture of multiple cells. In contrast, in situ sequencing [45, 21] and in situ hybridization based SRT such as 10x Genomics Xenium [19, 32], merFISH [9], CosMx (NanoString), MERSCOPE (Vizgen) measure the expression for a subset of pre-selected genes at cellular or sub-cellular resolution. The number of mRNA molecules measured for each gene varies considerably across different platforms.

Given the varying properties of individual SRT platforms, it is advantageous to integrate information from two or more platforms; e.g. combining information from whole-transcriptome platforms with lower spatial resolution with platforms that measure expression of a limited number of genes at high spatial resolution. Such integration could assist in two tasks: (1) predicting the expression of genes that are missing in the high resolution SRT datasets by referencing its expression in the low resolution SRT dataset, a process known as imputation; (2) inferring the mixture proportion of different cell types in a spot from the low resolution SRT dataset by using the high resolution SRT dataset as a reference, commonly referred to as deconvolution.

Multiple methods have been introduced to impute gene expression across single-cell sequencing technologies including scRNA-seq and single-cell sequencing assay for transposase accessible chromatin (scATAC-seq) [22, 39, 28, 8, 12, 47]. These tools commonly integrate multiple single-cell modalities into a shared latent space, allowing them to impute gene expression in cells on one modality by identifying nearby cells from a complementary modality within the latent space. A common way of modeling the latent space is by finding a low-dimensional factorization of scRNA-seq gene expression matrices [25, 49, 3, 4, 34, 42].

Similarly, in the spatial transcriptomics domain the SRT gene expression is traditionally imputed by integrating with a reference scRNA-seq dataset, often disregarding the spatial information inherent in the SRT dataset, treating it as a non-spatial modality [27]. Other methods such as Tangram [5], SpaGE [1] and SpaOTsc [7] imputes genes in SRT dataset by learning a mapping between each spatial location and the reference single-cells. On the other hand, akin to the modeling of single-cell datasets, several recent methods use low-dimensional factorization [10, 41] to model SRT data taking into account the spatial information. However, these methods are only applicable to a single dataset, and therefore not suitable for imputation across multiple SRT modalities.

All of the above mentioned tools used for imputation implicitly assume at least one of the modalities is scRNA-seq. As a result, applying them to impute gene expression across two SRT datasets faces two significant limitations. First, the spatial information in the datasets is ignored by the algorithm, and second, the reference dataset is assumed to have single-cell resolution. These limitations have been partially addressed by recently developed spatial alignment based tools, SLAT [48], and SANTO [24] which impute missing gene expression in one of the SRT datasets based on a learned spatial alignment. These tools do not handle platform-specific differences and overlook the need of deconvolution in case of low spatial resolution. To address the issue of admixed gene expression in technologies with multicellular spatial resolution, a number of methods have been developed to deconvolve ad-mixed SRT spots [2, 30, 44]. But these tools are not designed for imputation across two SRT datasets and furthermore they assume perfectly matched gene sets across the datasets, making them unusable for imputing gene expression in one SRT data from another.

We introduce **S**patial **I**ntegration for **I**mputation and **D**econvolution (SIID), an algorithm that reconstructs a latent gene expression matrix, jointly solving the imputation and the deconvolution problems. We employ a non-negative matrix factorization (NMF) based approach that takes advantage of the spatial structures of both SRT datasets, namely, superior resolution of one (e.g. 10x Xenium), and the full gene-panel present in the other (e.g. 10x Visium). We achieve this simultaneous deconvolution and imputation by inferring a latent gene expression matrix that has single-cell resolution but measures the whole transcriptome. From this matrix both Xenium and Visium expressions are generated where the Visium expression is obtained by mapping Xenium cells to Visium spots in physical space, and assuming the latent gene expression matrix admits a low-rank NMF. Intuitively this low-rank decomposition is representative of underlying cell types. Similar to existing NMF based methods, SIID constructs a lower dimensional latent gene expression matrix, but is different in several key aspects: First, we explicitly model counts as Poisson samples. Second, most existing methods either work with a single modality, or fit two NMFs sharing one of the component. In our model, the NMFs of two SRT datasets share both components via spatial mapping. Third, by sharing both NMF components, our model learns the cell type assignments for ad-mixed spots in a way that respects spatial information.

**Figure 1:**
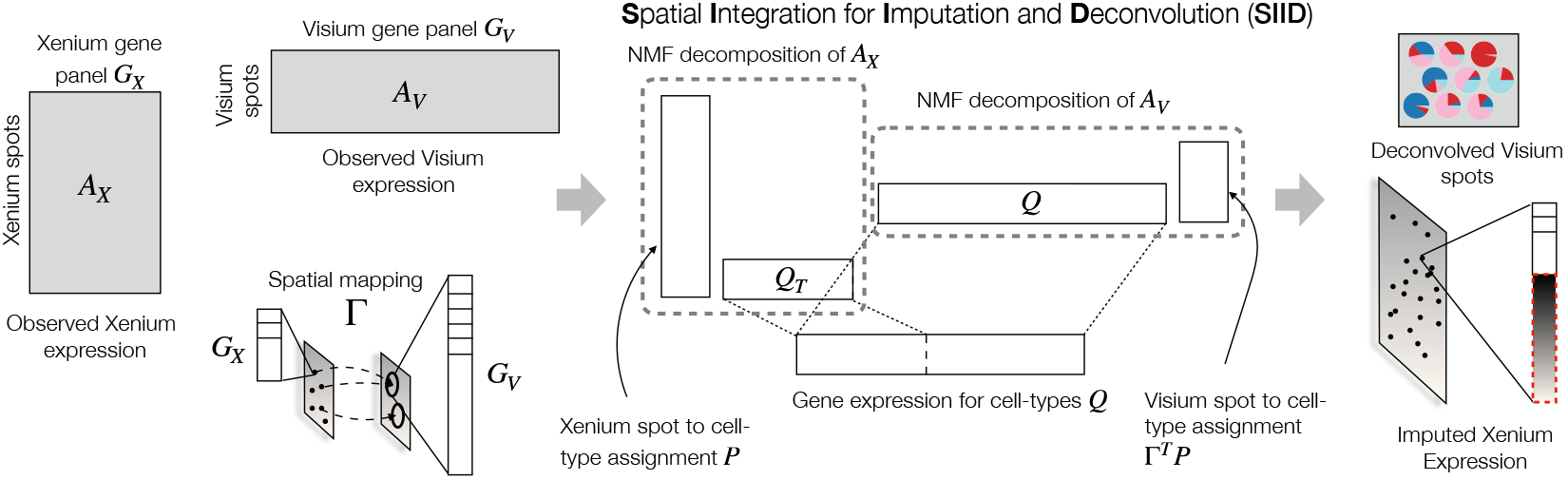
Overview of SIID. Given Xenium and Visium expression matrices *A*_*X*_ and *A*_*V*_, respectively, with corresponding gene panels *G*_*X*_ and *G*_*V*_ (with *G*_*X*_ ⊂ *G*_*V*_), and a Xenium to Visium spatial mapping Γ, SIID finds a NMF decomposition of a latent gene count matrix *A*_*U*_, with corresponding factorizations of *A*_*X*_ and *A*_*V*_, into location to latent factor assignment matrix *P* and the gene expression matrix *Q* for latent factors. The estimated parameters are used to impute the absent gene expression in Xenium data and predict the cell type mixture proportions for each Visium spot.

We demonstrate the effectiveness of SIID in both simulated and real data by evaluating the imputed gene expression and the inferred cell type mixtures. In simulated data, our method simultaneously deconvolves and imputes a paired dataset with highly ad-mixed Visium spots, which are intrinsically hard for non-spatially-aware methods. We benchmark imputation of Xenium genes through two paired Visium-Xenium datasets on breast cancer and colorectal cancer by holdout experiments and inspecting imputation of out-of-panel genes. SIID achieves highest performance in imputation measured by average holdout *R*^2^ score across entire Xenium gene panel, recovers most annotated cell types, and produces reasonable cell type deconvolution.

## 2 Methods

### 2.1 Representing paired spatially resolved transcriptiomics data

We represent a spatially resolved transcriptomics (SRT) dataset by a pair (*A, S*) where *A* ∈ ℕ^|*S* |×|*G* |^ is a gene count matrix and *S* ∈ ℝ^|*S* |×2^ is the two-dimensional physical coordinates for each spatial location, where *G* represents the set of genes measured in the SRT experiment. Suppose we are given two SRT tissue slices (*A*_*X*_, *S*_*X*_) and (*A*_*V*_, *S*_*V*_) from the same tissue measured with two different protocols which we denote by *X* and *V*. We assume that *X* has higher spatial resolution and a limited set of genes (small-panel), while *V* has lower spatial resolution but contains a superset of measured genes (large-panel), meaning |*S*_*X*_| ≫ |*S*_*V*_| and *G*_*X*_ ⊂ *G*_*V*_. In this manuscript *X* and *V* originate from 10x Xenium and 10x Visium platforms respectively.

We further assume that there exists an alignment between (*A*_*X*_, *S*_*X*_) and (*A*_*V*_, *S*_*V*_); i.e. there is a spot-spot correspondence between two slices as computed from any method that aligns two SRT datasets [50, 26, 40, 20, 11, 24]. Given the spot-spot correspondence, the coordinates *S*_*X*_ and *S*_*V*_ can be transformed to represent locations in a shared coordinate system between the SRT datasets. After such spatial transformation, the spot-to-spot correspondence matrix is represented with a binary mapping matrix 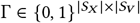, where Γ[*i, j*] = 1 if and only if the Xenium spot *i* is mapped to the Visium spot *j*. Our proposed framework works with any arbitrary Γ.

### 2.2 A shared cell type model between paired SRTs

We assume that *A*_*X*_ and *A*_*V*_ are sampled from a latent gene count matrix *A*_*U*_ based on the following assumptions.

- There exists a latent SRT dataset *U* represented with (*A*_*U*_, *S*_*X*_), where 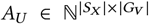 is the latent gene expression matrix. By construction *U* has same spatial coordinates as *X* and the same gene set as *V*.
- *A*_*U*_ follows a Poisson distribution with mean 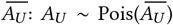. As *A*_*X*_ is a submatrix of *A*_*U*_, it naturally follows that 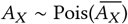 where 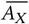 is a submatrix of 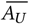 with columns restricted to *G*_*X*_.
- *A*_*V*_ follows a Poisson distribution with mean 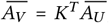, where 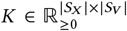 and *K* ° (1 − Γ) = 0, where °represents Hadamard product. *K*[*i, j*] represents the weight of contribution of a Xenium spot *i* to construct the Visium spot *j*, and is zero wherever Γ[*i, j*] = 0.
- 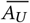 has a low-dimensional non-negative matrix factorization (NMF), which we denote as 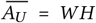, with 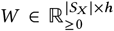 and 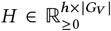, where *h* is the number of latent factors. This implies *A*_*X*_ has Poisson mean 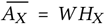 where *H*_*X*_ is a submatrix of *H* with columns restricted to *G*_*X*_, and *A*_*V*_ has Poisson mean 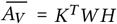.

With these assumptions, we formally state our inference problem as follows.

#### Paired NMF Inference Problem

Given a pair of SRT slices 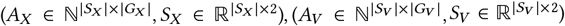 and spatial mapping matrix 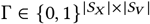, find nonnegative matrices 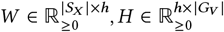, and 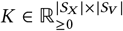 that solve the following problem:

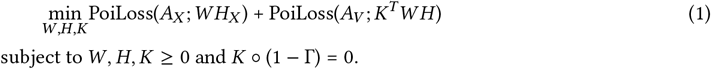

Here *K* corresponds to mixture weights matrix, where *K*[*i, j*] represents the contribution of Xenium spot *i* to the Visium spot *j* and PoiLoss(*Y*; *Z*) = (*Z* − *Y* log *Z*) is the negative log-likelihood for observing *Y* ~ Pois(*Z*).

#### Count-scaled Reparameterization

For numerical stability and better interpretation, we solve a reparameterized version of the Paired NMF Inference Problem. Recall 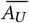 has a low-dimensional NMF 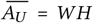, from which we derive estimated expression 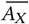 and 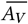. We rewrite this factorization with an alternative formulation where 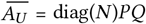 and where:

- 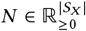 is a vector where *N* [*i*] is the inferred total gene counts for spot *i* in *U*, and 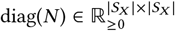 is a matrix whose diagonal elements are *N*,
- 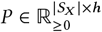, where each row *P* [*i*] is the normalized latent factor composition of spot *i* in *U*, i.e. 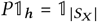,
- 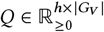, where each row *Q*[*j*] is the normalized expression of latent factor *j* in *G*_*V*_, i.e. 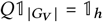.

We first show the reparameterized problem is equivalent to the original.

##### Lemma 1.

Given 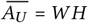 there exists *P, Q, N* as defined above such that 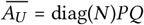, and vice versa.

*Proof*. see Appendix A.

From 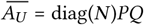 we have 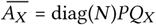, where *Q*_*X*_ is a submatrix of *Q* with columns restricted to *G*_*X*_. We next reparameterize *K*, the mixture weight. We define *M* = diag(*N*)*K*, and thus 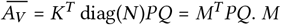 replaces *K* as the variable for inference and has identical constraints of *K*: *M* ≥ 0, *M* ° (1 − Γ) = 0. The reparameterized version is formally stated as follows:

#### Reparameterized Paired NMF Inference

Given a pair of SRT slices 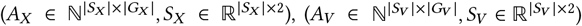 and spatial mapping matrix 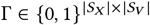, find nonnegative matrices 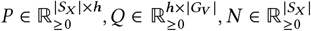, and 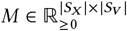 that solve the following problem:

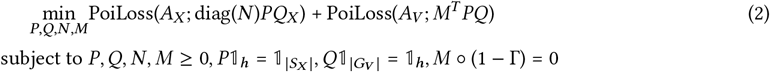

*M* corresponds to the mixture weights with *M*[*i, j*] being the contribution of gene counts from Xenium spot *i* to Visium spot *j*, and as above PoiLoss(*Y*; *Z*) = (*Z* − *Y* log *Z*) is the negative log-likelihood for observing *Y* ~ Pois(*Z*).

The reparameterized problem allows easier interpretation of the inferred model.

### 2.3 Implementation, parameter inference and imputing missing genes

We implement SIID to solve the Reparameterized Paired NMF Inference problem (Equation (2)). SIID solves the optimization problem in PyTorch ^1^ using gradient descent to optimize the model parameters. We use Adam optimizer and train for 5000 epochs, with a learning rate of 0.05.

#### Parameters of the model and 𝓁_2_ regularization

Since *P* and *Q* are row-normalized, we represent them as softmaxed matrices. Similarly, *N, M* are represented as exponentiated matrices. For numerical stability reasons, we place a 𝓁_2_ regularization with weight 10^−5^ on the parameters of the model (*P, Q* before softmax, *N, M* before exponentiation) [5].

#### Platform normalization

Our model assumes existence of a latent gene expression matrix *A*_*U*_ that in turn generates observations *A*_*X*_ and *A*_*V*_. In practice, if the two SRT datasets are generated from different SRT platforms, we expect platform-specific effects that should be modeled on top of the latent gene expression. Observing that the same gene could be expressed at different rates in different SRT datasets [6], we introduce a gene-wise scaling 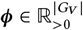, as a multiplier to the columns (corresponding to genes) of 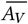 (to reduce non-identifiability in practice, we fix ***ϕ***[*j*] = 1 if *j* ∉ *G*_*X*_). More specifically, when platform normalization is enabled, the loss function of the Reparameterized Paired NMF Inference (Equation (2)) is updated to PoiLoss(*A*_*X*_; diag(*N*)*PQ*_*X*_) + PoiLoss(*A*_*V*_; *M*^*T*^ *PQ* diag(***ϕ***)).

#### Entropy regularization

To assign the spots in *X* into distinct cell types, we optionally add an entropy regularization term of ℋ = −*ω* (*P* log *P*) to the loss function (2) with increasing weight *ω* = exp(*k/λ*) across training epochs, where *k* is the current epoch and *λ* is a hyperparameter. Lower entropy encourages each Xenium spot to be predominantly assigned to a single latent factor or cell type, rather than being evenly distributed across multiple types. In general, when the focus is imputing missing genes, we suggest a larger value of *λ* such that the Poisson losses are the dominant terms. When the focus is the deconvolution of admixed spots, we suggest a smaller value of *λ* such that entropy regularization becomes more dominate near the end of the training process. We use *λ* = 500 in simulation and deconvolution on real data (Section 3.2, Section 3.4) and *λ* = 1000 for imputation on real data (Section 3.3).

#### Random restarts

As NMFs are known to be sensitive to initializations [15, 16], we always restart our model 3 times and use the one with best loss value.

#### Imputing missing genes

To impute the expression of a missing or holdout gene *g* on *X*, the model is trained with *g* ∉ *G*_*X*_ and *g* ∈ *G*_*V*_. With trained parameters *P, Q* and *N*, the imputed expression for *g* is diag(*N*)*PQ*_*g*_ where 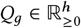 is the column of *Q* corresponding to *g*. Multiple genes can be imputed in the same run.

## 3 Results

### 3.1 Setup and evaluation

For simulation and evaluation, we use two publicly available datasets of paired Xenium and Visium SRT from adjacent tissue sections: one from breast cancer (BRCA) pair [19] and another from colorectal cancer (CRC) pair [32]. Xenium, Visium and scRNA-seq data for both datasets (and cell type annotation for BRCA) are downloaded as described in Appendix C.1. Furthermore, Appendix C.2 contains detailed dataset statistics.

#### BRCA pair

We align Xenium and Visium slices by first determining the correct physical scale, manually identify key points between slices, then using key point registration to fit a rigid transformation mapping the Visium coordinates to Xenium [17]. With mapped coordinates, Γ is generated by matching each Xenium spot to its closest Visium spot up to a distance of 100 *µ*m.

#### CRC pair

We align Xenium and Visium slices by first determining the correct physical scale, then instead of manual key point registration, we run PASTE2 [26] and compute a rigid transformation from the PASTE2 outputs to map Visium coordinates to Xenium. Γ is generated by matching each Xenium spot to its closest Visium spot up to a distance of 100 *µ*m.

#### 3.1.1 Imputation setup and holdout evaluation

For both the BRCA and CRC datasets, we divide Xenium panel genes into 10 equal-sized folds. For each fold, we remove the expression of genes in fold from the Xenium dataset, then run the imputation algorithm to obtain estimates of these holdout genes. To evaluate the imputation results, for each gene in the fold, we compute the *R*^2^ score between estimated expression for each Xenium cell and the observed counts (which were held out during training). The *R*^2^ score between two vectors (*x, y*), also called coefficient of determination, equals 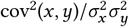 where cov is the covariance and *σ*^2^ is the variance. The *R*^2^ score for a given model is the average gene-wise holdout *R*^2^ across all 10 folds, where each gene is a holdout exactly once across 10 folds.

### 3.2 SIID recovers the cell type expression in simulated dataset

We evaluate the performance of SIID on simulated paired Xenium and Visium dataset generated from the BRCA dataset [19]. The evaluation focuses on three aspects: imputation of genes not present in the Xenium data, recovery of the spatial location to cell type assignments, and accuracy of cell type gene expression profiles.

To simulate a realistic expression profile, we use the gene expression for the scRNA-seq dataset and the spatial coordinates from the Xenium and Visium data. For this simulation, we used clusters (see Appendix B.1) obtained from unsupervised clustering of the scRNA-seq data and define them as cell types. Each Xenium spot is assigned to a cell type using a checkerboard spatial pattern (as described in [18]) (Figure 2A), where each grid on the checkerboard has a different cell type from its neighbors. We use the average gene expression for individual cell type from the scRNA-seq dataset to simulate the gene counts for Xenium spots according to their cell type. Expression data for Visium spots are generated by applying the spatial mapping from Xenium to Visium data (see detailed steps in Appendix B.1).

**Figure 2:**
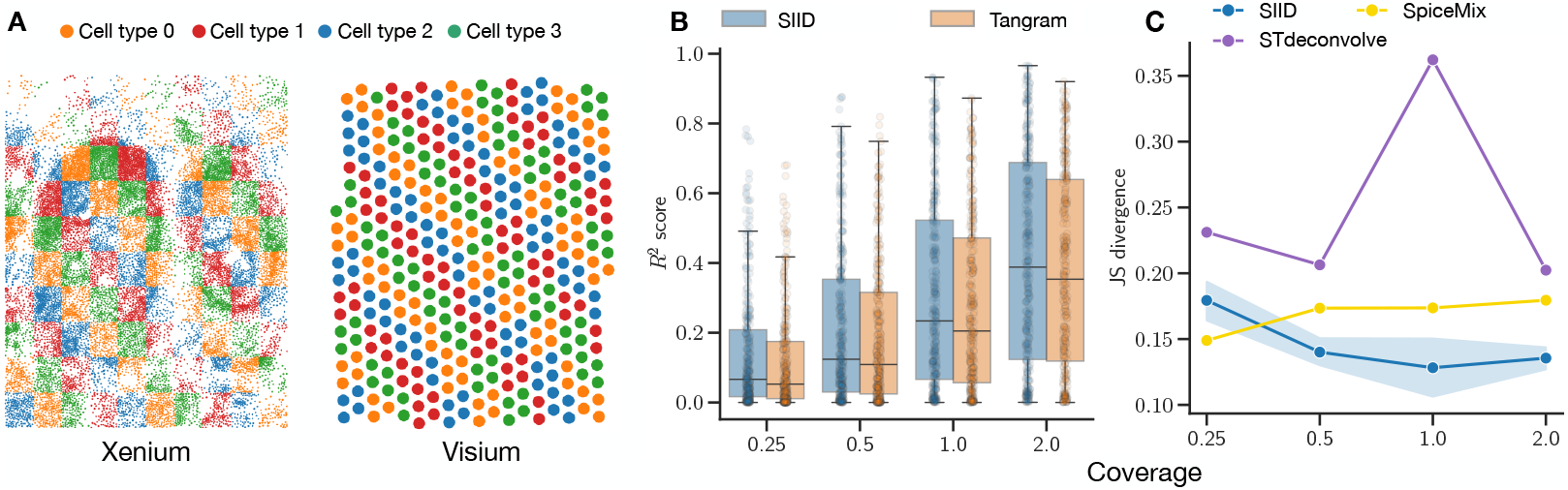
**A**: A simulated Xenium and corresponding Visium dataset with gene expression of cell types obtained from a matching scRNA-seq dataset [19]. **B**: *R*^2^ of imputed gene expression for holdout genes from SIID and Tangram stratified by coverage. **C**: Average Jensen-Shannon (JS) divergence between ground truth and predicted cell type mixture proportions predicted by SIID, STdeconvolve, and SpiceMix for simulated Visium data over all spots stratified by coverage.

We created a variety of simulation datasets by varying i) the number of grids, *l* on each side of the checkerboard with *l* ∈ {10, 20} ii) gene counts generated for each Xenium spots, iii) the number of cell types *h*, where *h* ∈ {4, 8, 16}. To simulate Xenium gene expression we began with the actual vector of UMI counts *N*_*X*_ from BRCA Xenium dataset. This was then scaled using a coverage fraction *ρ* (which we refer as *coverage*) to create a UMI count vector 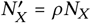, with *ρ* ∈ {0.25, 0.5, 1, 2} which is used for simulating gene expression for Xenium spots. By varying *l, h, ρ*, we created in total 24 distinct configurations each representing varying levels of difficulty. We also study the effect of additional factors in simulation, namely, spatially variable coverage, noise in spatial mapping matrix, and gene-wise scaling factor in the imputation results (see Appendix B.3, B.5, and B.6).

We evaluate the accuracy of SIID for gene expression imputation and cell type deconvolution in simulations, comparing its performance to Tangram and STdeconvolve. For the configuration with *l* = 20 grids and *h* = 8 cell types, SIID achieves superior *R*^2^ scores when compared to Tangram [5] across different coverage levels (Figure 2B). On the same simulation configuration, SIID outperforms STdeconvolve, and SpiceMix by achieving the lowest average Jensen-Shanon (JS) divergence (Figure 2C) between the predicted cell type mixture proportion (see Appendix B.2.3) and the ground truth. On other simulation configurations (see Figure S1), SIID consistently receives the best *R*^2^ scores when compared to Tangram and comparable JS divergence when compared to STdeconvolve, and SpiceMix (see Appendix B.2).

### 3.3 Imputing missing genes in real data

To perform benchmarking on real data (see Section 3.1.1), we run SIID with number of latent factors *h* = 20 for BRCA and 40 for CRC datasets, entropy regularization with *λ* = 1000, platform scaling enabled, 5, 000 training epochs and 3 restarts. For training we use genes that are available in both Xenium and Visium datasets, excluding the ones in holdout set. We document the details of hyperparameters in Appendix C.3.

We benchmark four existing methods [23] for evaluating the imputation results - Tangram [5] (in cell and cluster modes), gimVI [27], SLAT [48] and SANTO [24] (with pre-computed mapping Γ). However, we do not present results for gimVI [27] due to runtime errors. In addition, we present four baseline approaches by using the spatial mapping Γ; Baseline A imputes gene expression for each Xenium cell by simply using the gene expression from the corresponding Visium spot based on the spatial mapping, Baseline B is a variant of Baseline A that takes total count per Xenium cell into account, and Baselines C and D employ *k*-nearest-neighbor based smoothing based on Baselines *A* and *B*, respectively. Detailed procedures for benchmarking are in Appendix C.4, and runtimes for benchmarking are in Appendix C.6.

#### SIID accurately imputes genes in the holdout experiments

Compared to other methods, SIID achieves the best *R*^2^ scores in both datasets - BRCA and CRC on average holdout *R*^2^ scores across all ten folds (Table 1). Among the competing methods, Tangram performs reasonably well when run in cell mode (referred as Tangram (cell) in Table 1). In addition to *R*^2^ scores, SIID also consistently outperforms other benchmarked methods on four other correlation metrics ([23], Appendix D.1). Additionally, SIID continues to show robust performance in the presence of noise in spatial mapping matrix (Appendix C.5).

**Table 1:** Comparison of average holdout *R*^2^ scores across methods. Bold indicate best performer.

| Dataset | SIID | Tangram |  | SLAT | SANTO | Baselines |  |  |  |
| --- | --- | --- | --- | --- | --- | --- | --- | --- | --- |
|  |  | cell mode | cluster mode |  |  | A | B | C | D |
| BRCA | **0.2527** | 0.1874 | 0.1556 | 0.0688 | 0.1042 | 0.0373 | 0.0807 | 0.0766 | 0.0664 |
| CRC | **0.2248** | 0.1789 | 0.1382 | 0.0608 | 0.0727 | 0.0325 | 0.0573 | 0.0443 | 0.0412 |

**Table 2:** Comparison of average holdout R2 scores using scRNA-Xenium pairing.

| Dataset | SIID (Visium) | Tangram (scRNA, cell) | Tangram (scRNA, cluster) |
| --- | --- | --- | --- |
| BRCA | **0.2527** | 0.2195 | 0.2203 |
| CRC | **0.2248** | 0.1960 | 0.1822 |

SIID also achieves a higher *R*^2^ score for most of the individual genes compared to the closest competitor (Figure 3A). For this evaluation, we compare SIID with Tangram (in cell mode) by evaluating the *R*^2^ scores of both methods for each gene in the Xenium panel from the BRCA dataset (see Figure S5A in Appendix D.2 for CRC) along with the total UMI counts. We observe SIID outperforms Tangram overall, while Tangram performs better for a small number of genes, most of them lowly-expressed. We further visualize the difference of *R*^2^ scores and its relationship to expression level in Figure S6. As SIID is a generative model, we can also validate the model by showing its ability to predict the sparsity of holdout genes (Figure S7).

**Figure 3:**
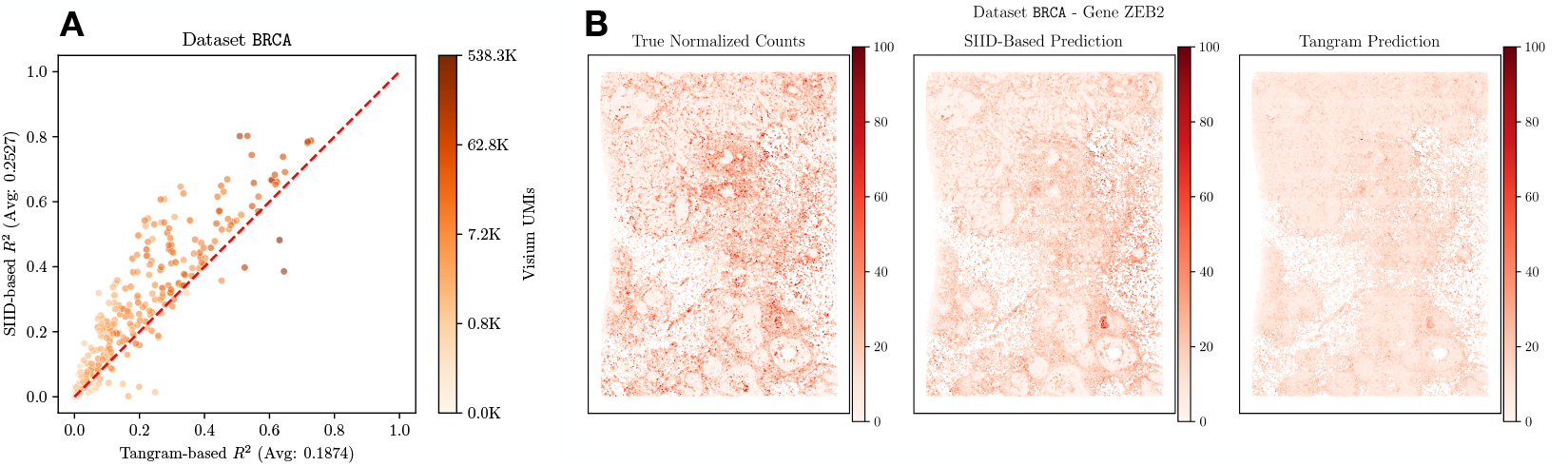
**A:** Comparison of holdout *R*^2^ score of SIID (Y-axis, correlation score averaged over 5 runs) and Tangram in cell mode (X-axis) for each gene in the Xenium panel for the BRCA dataset. Each point on the plot corresponds to a single gene, whose color corresponds to the number of Visium UMIs mapped to the gene on a log-scale. Genes above the red *y* = *x* line have higher imputation performance for SIID compared to Tangram. **B:** Visualizing expression of gene *ZEB2*: Left: Observed Xenium expression (ground truth for evaluating imputation), Middle: SIID prediction when the gene is held out (*R*^2^ = 0.4781 against ground truth), Right: Tangram prediction when the gene is held out(*R*^2^ = 0.2931 against ground truth). *R*^2^ between predicted expression and observed expression are in title of each plot. We normalize the total counts within each gene to 1, 000, 000 and plot all normalized counts on the same color scale.

#### SIID recovers spatial pattern of Xenium gene expression

SIID is superior to Tangram in recovering spatial patterns for certain marker genes [19] (see Figure 3B and Figure S9). In particular, we found that the SIID-imputed expression for *ZEB2*, a well-known oncogenic driver implicated in epithelial-mesenchymal transition (EMT) in breast cancer [14, 13], closely mirrors the true tissue structure (*R*^2^ = 0.48), while Tangram’s imputed expression is more uniformly distributed across the slice (*R*^2^ = 0.29, Figure 3B). We present similar results for other marker genes in Appendix D.2, Figure S9.

#### Comparison to imputing with paired scRNA-seq

On imputing Xenium genes, SIID with Visium data outperforms Tangram paired with a scRNA-seq reference^2^ for both modes and both datasets (Table S4). SIID is not specifically designed to impute Xenium gene expression with a non-spatial reference, but Tangram can be run in both cell and cluster mode for this setup. In order to run Tangram on the CRC dataset in a reasonable time, we down-sampled the dataset as described in Appendix C.4. To emphasize, despite not using the high resolution scRNA-seq data, SIID outperforms Tangram with access to scRNA-seq data, although with a smaller lead.

### 3.4 Deconvolving cell types in real data

We also evaluate SIID’s performance in deconvolving the admixed gene expression in Visium data using the paired Xenium data. For deconvolving Visium spots without annotation, we run the same setup as described in Section 3.3 and Appendix C.3, with no holdout genes and *λ* = 500 for entropy regularization. Here we evaluate only with the BRCA dataset, as this dataset includes annotated cell types for both the Xenium and scRNA-seq data provided by 10x Genomics.

#### SIID recovers most annotated major cell types

SIID recovers most annotated cell types with more than 1000 cells by mapping them to one or a few latent factors (Figure 4A). To benchmark this, we train the model on the BRCA dataset. Since the reference annotation assigns each cell to exactly one type, we similarly assign each Xenium cell to one of 20 latent factors by finding which latent factor contributes the most to its inferred expression, i.e. Xenium cell *i* is assigned to latent factor arg max_*j*_ *P* [*i, j*]. We calculate the cosine similarity (see Figure 4A) of this clustering of cells to the cell type annotation provided by 10x Genomics, which contain 19 distinct cell types (besides “Unlabeled”) and some annotations are more granular than others. In addition, we find that the clustering of Xenium cells by SIID is more similar to the 10x Genomics annotation (Adjusted Rand Index (ARI) =0.460) than the Leiden clustering (with 21 clusters) is to the 10x annotations (ARI = 0.347).

**Figure 4:**
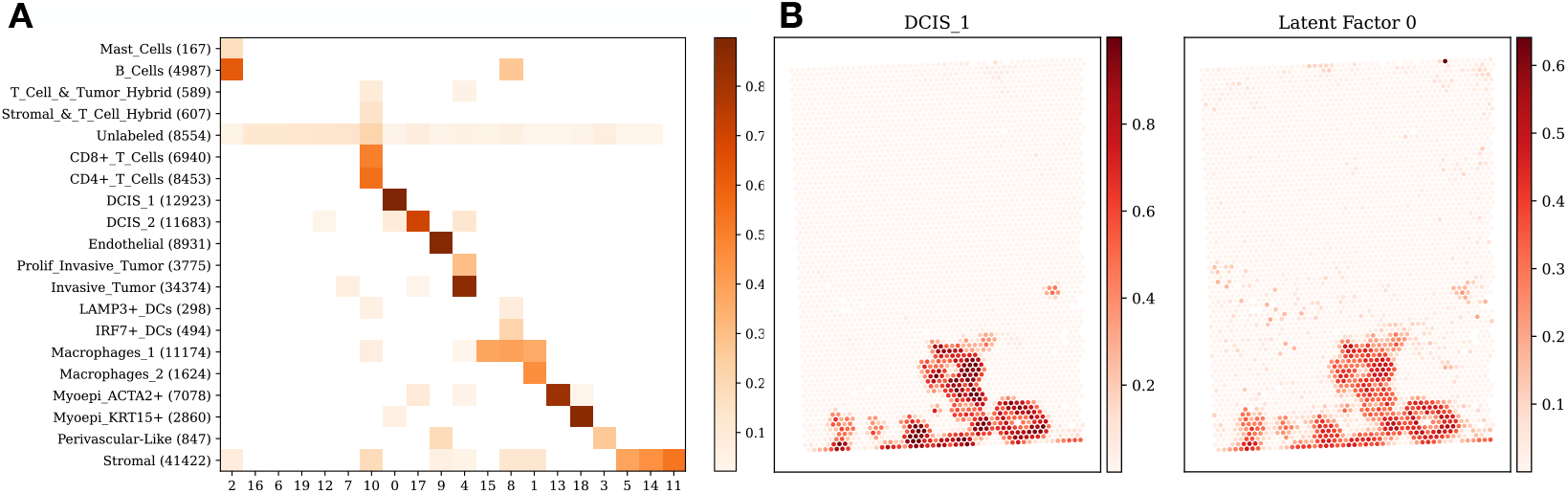
**A:** Comparison of the clustering of Xenium cells obtained by SIID on the BRCA dataset with the cell type annotation provided by 10x Genomics. Each row corresponds to a annotated cell type, with the number of Xenium cells belonging to that type in parentheses. Each column corresponds to a latent factor derived by SIID. Intensity of each grid point is the cosine similarity between the assignments, with similarity scores below 0.02 excluded for visual clarity. **B:** Spatial distribution of deconvolved DCIS 1 cell type by RCTD and one of the latent factors of SIID.

#### SIID recovers spatial distribution of deconvolved cell types

SIID’s unsupervised deconvolution of Visium spots are visually similar to the supervised deconvolution performed by RCTD [6], a leading method for deconvolving admixed SRT data (Figure 4B). For SIID, we train the model on the BRCA dataset (with Xenium and Visium data), and obtain the unsupervised latent factor deconvolution for each Visium spot by computing *M*^*T*^ *P* and normalizing by each spot. We run RCTD [6] with Visium and scRNA-seq data annotated with cell types to obtain a supervised cell type deconvolution of Visium spots (cosine similarity plot in Appendix D.2, Figure S5B). We observe that the spatial distribution of deconvolved DCIS 1 cell type from RCTD and latent factor 0 from SIID have strong spatial agreement (Figure 4B). We emphasize that other imputation tools such as Tangram are not capable of determining cell types for spatial location.

## 4 Discussion

SRT technologies are constantly evolving, with new techniques for measuring gene expression in physical space emerging in different flavors. Each SRT technique offers unique insights, but there are many practical considerations and trade-offs to keep in mind, such as spatial resolution, total observed counts, gene panel size, processing time, and costs. Selecting the most appropriate SRT protocol for profiling a target sample can be challenging, and it is often advantageous to adopt a multi-modal approach. Using multiple SRT methods to study the same tissue allows for complementary insights, combining high spatial resolution, adequate coverage (with high counts for each gene), and a comprehensive whole-transcriptomic view. This approach helps to mitigate the limitations of individual modalities, providing a more complete measurement. However, most existing methods for integrating SRT slices either focus on integrating without spatial information, essentially treating SRT datasets as scRNA-seq datasets, or are designed for integrating SRT slices from the same modality and largely ignore potential differences between modalities. Few existing methods consider alignment and integration of different SRT modalities and perform simultaneous imputation and deconvolution.

We introduce SIID, a new approach of integrating SRT datasets that simultaneously performs imputation on a high resolution targeted SRT dataset (Xenium in our experiments) and deconvolution on a low resolution ad-mixed SRT dataset (Visium in our experiments), using each other as the reference dataset instead of relying on a third scRNA-seq one. This task is achieved by our proposed joint NMF inference framework, where in addition to shared gene expression for each latent factors, the latent factor per spot are also shared up to a convolution operation. This allows us to recover the latent SRT dataset which simultaneously reveals the whole-transcriptomic imputation of the Xenium data, gene expression of the cell types, and deconvolution of the Visium spots into cell type proportions. We validate SIID on both simulated and real datasets by recovering the simulation configuration in a highly ad-mixed setup, and achieving higher holdout *R*^2^ scores on two different real dataset pairs.

There are several limitations of SIID which are directions for further improvement. First, the SIID model is capable of inferring whole-transcriptomic expression for 100, 000s of cells, thanks to the fact that SIID does not require explicit construction of the latent gene expression matrix 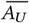. However, in current validation procedures, we do not use out-of-Xenium-panel genes that are present in Visium datasets, effectively discarding 95% of Visium counts (Appendix C.2). A potential reason for not fully utilizing these additional genes is that we currently account for platform specific differences solely through the platform scaling factors. More advanced models that account for the expression differences could enable better utilization of out-of-panel gene expression during inference. Second, our current method is reliant on an accurate spatial mapping matrix Γ, which is not always available. We plan to reduce reliance on the spatial mapping by using a more flexible Γ that maps each Xenium cell to multiple (or even all) Visium spots, taking advantage of a scRNA-seq reference when available, or integrating with an existing SRT alignment approach to refine the alignment using the inferred model. Third, our method is currently uses a Poisson count model, but can easily be extended to other probability distributions. We briefly discuss an alternative implementation where we model Visium counts using negative binomial distribution with gene-specific overdispersion in Appendix C.7. Finally, our model contains a number of hyperparameters that requires understanding of the underlying data. For example, the number of latent factors is a very important matter as it directly impacts interpretation and analysis of the inferred model. The choice of hyperparameter *λ* in the entropy regularization affects the number of training epochs and the balance between imputation and deconvolution. In future extensions, we will explore venues of automatically inferring these hyperparameters depending on the use case.

Recently, new SRT technologies such as Xenium 5K and Visium HD [32], capable of measuring a large panel of genes in sub-cellular resolution, have emerged. However, high resolution datasets are often significantly sparser than low resolution or small-panel datasets. Integration methods such as SIID can be adapted to improve coverage of these sparse datasets, allowing for accurate characterization of the target tissue. Overall, SIID is a robust and interpretable method that is scalable and flexible to handle new SRT technologies.

## Acknowledgments

This research is supported by National Cancer Institute (NCI) grants U24CA248453 and U24CA264027 to B.J.R and by the Princeton Ludwig Branch.

## Appendix

### A Proof of the Reparameterization Lemma

To formally prove Lemma 1 we require no empty rows (rows will all zeros) in *W* and *H*. This means there are no spots with zero inferred expression, and there are no latent factors with zero inferred expression.

*Lemma*. Given 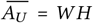 where *W* and *H* has no empty rows and *W, H* ≥ 0, there exists *P, Q, N* with 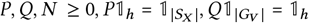 as defined in Section 2.2, such that 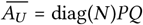, and vice versa.

*Proof*. The forward direction can be proved in two steps, starting with *W* and *H* :

- Let 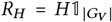 be the row sum of *H*. Since *H* has no empty rows, *R*_*H*_ *>* 0 and let *W*_1_ = *W* diag(*R*_*H*_), *Q* = diag(1*/R*_*H*_)*H*. We have *W H* = *W*_1_*Q* where Q is already row-normalized.
- Let *N* = *W*_1_𝟙_*h*_ be the row sum of *W*_1_. We can further set *P* = diag(1*/N*)*W*_1_ knowing *N >* 0, and since diag(*N*)*P* = *W*_1_, we have diag(*N*)*PQ* = *W*_1_*Q* = *W H*.

The reverse direction can be proved by setting *W* = diag(*N*)*P, H* = *Q* from which we get *W H* = diag(*N*)*PQ*.

### B Benchmarking on simulated Xenium and Visium data

#### B.1 Steps for generating simulated Xenium and Visium data

We collect the paired scRNA-seq data, Visium data, and Xenium data from the BRCA dataset [19] as described in Section 3.1. Following the notation of the Section 2.2 the simulated Xenium slice measures genes in *G*_*X*_ over cell locations *S*_*X*_, the simulated Visium slice measures genes in *G*_*V*_ over spot locations *S*_*V*_, and *G*_*X*_ ⊂ *G*_*V*_. We further cluster the cells in the scRNA-seq data and selected *h* distinct clusters. Our simulation process contains the following steps.

- **Step 1: Checkerboard style spatial clusters for Xenium spatial locations** We first assign each cell in *X* to one of *h* clusters. This assignment is done by dividing the coordinates of *S*_*X*_ into square grids, then assigning all cells in each grid to the same cluster such that that cells from adjacent grids belong to different clusters. Such assignment will create a checkerboard style [18] pattern based on *S*_*X*_. This spatial clustering increases the number of cells that are spatially neighboring yet belong to different clusters, and as a result most Visium spots contain a mixture of cell types. It has been observed that state-of-the-art spatial clustering algorithms struggle to identify the cluster identity accurately [18]. The spatial location to cluster assignment is represented as a matrix 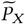, where 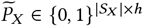.
- **Step 2: Constructing the Poisson mean for the latent gene expression matrix** Unlike real data, in simulation we can create the Poisson mean of the latent gene expression matrix 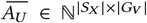. Given the scRNA-seq gene expression matrix and the cell to cluster assignments, we first compute the normalized average gene expression for each cluster, represented by 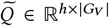 s.t. 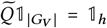. Using the Xenium spatial location to cluster assignment from step 1, the simulated latent gene expression matrix is sampled from a Poisson mean 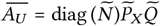, where 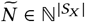 denotes the expected gene count for each Xenium cell.
- **Step 3: Simulating Xenium gene expression matrix** The Xenium expression matrix 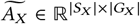 is a submatrix of the latent gene expression matrix 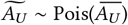 where columns are restricted to gene set *G*_*X*_.
- **Step 4: Simulating Visium gene expression matrix** We use a pre-computed mapping matrix Γ to compute the Poisson mean of Visium gene expression as 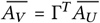, meaning the Poisson mean for each Visium spot equals the sum of Poisson mean from each Xenium cell mapped to that spot. We then sample 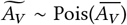 independently of 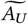. We additionally construct cluster mixture proportion matrix 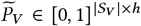 by row-normalizing 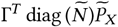.

Upon simulating the Xenium count matrix 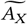 and Visium count matrix 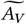, we can construct an instance 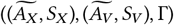 for *Reparameterized Paired NMF Inference* problem using the imputation setup described in Section 3.1.1 for one of the 10 folds. We then run SIID with *h* latent factors. We measure the accuracy of SIID, by comparing the estimated parameters 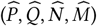 with the ground-truth parameters as follows,

- We compute Jensen-Shannon divergence between 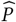 and 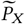 to measure the accuracy of Xenium spot to cluster assignment.
- To evaluate the cluster assignment for Visium spots, we first calculate 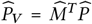 using estimates from SIID. The row-normalized 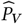 is compared with 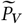 as defined above in Step 4 of the simulation.
- We evaluate the gene expression for inferred clusters by comparing 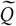 to 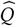. Here we compute the pairwise Pearson correlation coefficients between the rows of 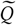, and 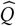.
- We also evaluate the performance of our model in imputing holdout genes in Xenium panel by the same process described in Section 3.1.1.

#### B.2 Benchmarking deconvolution results

##### B.2.1 STdeconvolve setup

To do a fair comparison with SIID, we run STdeconvolve on Visium data on the shared genes between Xenium and Visium data. As the remaining genes are low in number we did not perform any gene selection before running STdeconvolve. The other steps are same as described in STdeconvolve documentation ^3^.

##### B.2.2 SpiceMix setup

We follow instructions given in the SpiceMix paper [10] and SpiceMix GitHub repository^4^ and preprocess the Visium dataset as follows. First, all genes that are expressed in less than 10% of spots are removed. Second, all counts are log-normalized to a target UMI count of 10, 000 per cell. Third, the neighbourhood graph is derived by connecting each spot with its 6 closest neighbors. Afther these transformations, SpiceMix is run with the default parameters for 200 epochs, with the number of metagenes set to the number of hidden dimensions used in SIID and STdeconvolve.

##### B.2.3 Computing cell type proportion matrix

We first map latent factors inferred from SIID to true cell types gene expressions to find the best mapping between the latent factors and the cell types by a greedy match process. Given that correspondence we rearrange the columns of mixture proportion matrix *P* to match the true cell type proportions. A similar approach is used to find the best order for cluster mixture proportions from STdeconvolve. Finally, we use Jensen-Shannon divergence [29] (JS divergence) to evaluate the quality of predicted cell type mixture proportions when compared to the true mixture proportions. A lower JS divergence signifies that the predicted mixture proportion is closer to the ground truth.

#### B.3 Evaluating the effect of noise in spatial mapping

We evaluate the reliability of spatial mapping in the resulting imputation and deconvolution performance by introducing noise to the mapping matrix Γ. To control the level of noise, we randomly select *τ*% of existing mapping, i.e. (*i, j*) where Γ[*i, j*] = 1, and swap with a Xenium-to-Visium spot pair (*i*^′^, *j* ^′^) such that Γ(*i*^′^, *j* ^′^) = 0, resulting in the noisy spatial mapping matrix Γ^′^ has Γ^′^[*i*^′^, *j* ^′^] = 1, and Γ^′^[*i, j*] = 0. In simulation we vary *τ* ∈ {0, 10, … 100} denoting Γ^′^ = Γ, to Γ^′^ where all the spatial mappings are swapped. We observe SIID yields comparable *R*^2^ scores with different levels of noise introduced to Γ (see Figure S2). However the JS divergence, which measures the deconvolution performance for the Visium spots, increases as noise in the spatial mapping rises, indicating poorer deconvolution performance as the spatial mapping matrix becomes less reliable.

#### B.4 Execution time and memory usage

We measure the time and memory usage for running SIID and Tangram by profiling the execution of both the tools using Python’s `time` and `tracemalloc` modules. We note that the recorded time and peak memory usage for different simulation configurations remain relatively stable across different levels of coverage, grid sizes, and cluster counts, indicating that their computational costs are scalable across these parameters. While SIID requires more execution time than Tangram, Tangram requires significantly higher memory (see Figure S4).

#### B.5 Evaluation of spatially variable scaling factor

We considered a spatially variable coverage *ρ* (see Section 3.2) in contrast to a fixed value and evaluated the imputation performance of SIID on a simulated dataset (with *l* = 10, *h* = 8). The resulting simulation has a spatially changing scaling factor across the Xenium spots (see Figure S3A). The overall *R*^2^ score for the spatially variable scaling factor is slightly higher (see Figure S3 B) than the *R*^2^ score attained with a constant scaling factor (*ρ* = 0.5).

#### B.6 Evaluation of gene-wise scaling factor

We evaluate the effect of gene-wise scaling in the performance of SIID and Tangram, we created a simulation that uses gene-wise scaling while simulating the Xenium and Visium datasets. To emulate the real data, we first measure the gene-wise scaling factors for the shared genes between the BRCA Xenium and Visium data (see Figure S8 A). Later, we sample from this empirical distribution of gene-wise scaling factors to simulate the gene expression of the Visium data. While running SIID, we enable the platform normalization (see Section 2.3). We observe SIID achieves a superior *R*^2^ score (see Figure S8 B) as compared to Tangram in imputing gene expression for the holdout genes in Xenium dataset.

### C Detailed benchmarking information for real data

In this section, we describe detailed setups for running our method and other benchmarked methods over real data.

#### C.1 Downloading and preprocessing datasets

##### BRCA pair

We downloaded the following datasets from 10x Genomics website^5^: Xenium In Situ Sample 1, Replicate 1; Visium Spatial; FRP (Fixed RNA Profiling, as the scRNA-seq pairing). We also download the cell type annotation for FRP and Xenium in the same page.

##### CRC pair

We downloaded the following datasets from 10x Genomics website^6^: Visium CytAssist V2, Sample P2 CRC; Xenium In Situ, Sample P2 CRC; and Chromium Single Cell Flex, aggregated (as the scRNA-seq pairing). The aggregated Chromium dataset contains scRNA-seq data from 8 different samples, and we subset the it to only cells collected from the matching Sample P2 CRC (indices 1, 3, 5 and 7).

#### C.2 Dataset statistics

Table S1 contains detailed statistics regarding the data used for benchmarking. The rows indicated as (in-Xenium-panel) are the statistics for Visium dataset when restricted to paired Xenium panel.

#### C.3 Hyperparameters of SIID

SIID contains a number of hyperparameters. Most of these parameters are chosen based on common sense and a small-scale parameter search.

**Table S1:**
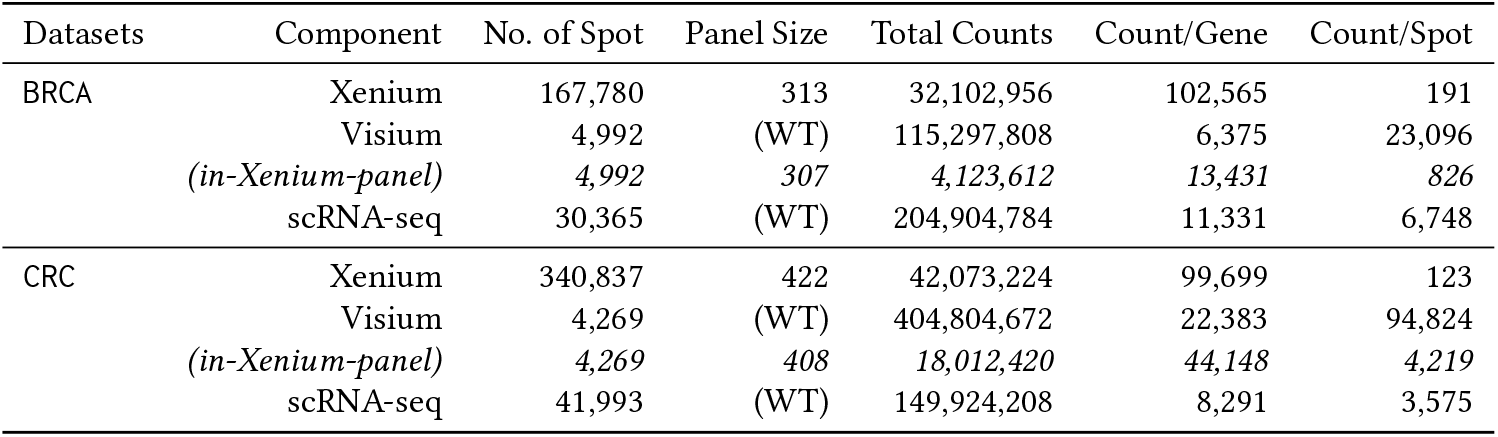
Dataset statistics. *(in-Xenium-panel)* denotes Visium dataset subsetted to matching Xenium gene panel. WT stands for whole-transcriptomic (18 thousand genes).

- Number of latent factors *h* determines the size of *P* and *Q* of the factorization, and is loosely based on how many cell types the datasets have. In simulation, *h* is set to match the data generation process, and in real data, it is set to 20 or 40 for `BRCA` and `CRC` respectively. This is the only dataset-specific parameter we employ.
- We described platform scaling technique in Section 2.3 as a way to model platform-specific count modeling between Xenium and Visium datasets. This is disabled for simulation both in dataset generation and inference, and is enabled when running with real data.
- SIID uses entropy regularization (Section 2.3) to promote assigning Xenium cells to smaller number of latent factors. With fixed total of 5,000 training epochs and regularization weight exp(*k/λ*) where *n*_epoch_ is the epoch number, higher *λ* means weaker regularization strength. We use *λ* = 1000 for real data imputation (Section 3.3), and *λ* = 500 elsewhere.
- SIID has the flexibility of using genes inside and outside the targeted panel during inference. For our runs, we choose to not use Visium genes outside Xenium panel (except those in holdout set for imputation).
- SIID is run with 3 restarts, and the run with lowest total loss is used for prediction. It also uses a 𝓁_2_ regularization over raw model parameters (Section 2.3), whose weight is set to 10^−5^ for all runs. The model is optimized with Adam optimizer with learning rate of 0.05.

#### C.4 Setup for benchmarking imputation

We select the methods to benchmark by the top performers in [23], excluding SpaOTsc [7] as it requires a pairwise distance matrix over Xenium data with over 100K cells, and excluding SpaGE [1] as it operates over log-normalized data while we benchmark imputation with raw counts. We additionally select SLAT [48] and SANTO [24] as representatives of SRT-alignment-based imputation methods.

- Tangram [5] is run with Visium dataset as the single-cell data, and the Xenium dataset as the spatial data with default parameter and 1000 training epoches. We also run Tangram in two configurations, once in default mode (cell mode), and once in cluster mode with cluster labels generated by running Leiden clustering [43] in scanpy package [46] with default parameters. Tangram with scRNA-seq pairing is run in the same way, except that we downsample the scRNA-seq dataset for CRC by 2× when Tangram is run in cell mode. This is so it could finish in a reasonable amount of time.
- gimVI [27] is run in default parameters with the same number of latent factors as SIID, and optimized for 200 epochs. As advised by authors, We also remove empty cell or spots before optimization. However, we could not finish optimization due to model parameters becoming NaN frequently in the middle of the optimization after a large number of attempts and trying over many different configurations.
- SLAT [48] is run with parameters recommended for Xenium-Visium pairing (120 neighbors for building kNN graph for Xenium, 5 neighbors for building kNN graph for Visium). For each Xenium cell (even outside of Visium-Xenium overlap area), SLAT matches it to a Visium spot. Imputation of a holdout gene is performed by copying the expression of such holdout gene on matched Visium spot to the Xenium cells.
- SANTO [24] proposes an imputation strategy from two aligned SRT slices based on Poisson regression, which we re-implement and benchmark. For each holdout gene *g*, we train a Poisson regression model of form Γ_*I*_*V*_*g*_ ~ Pois(*A*_*XI*_ *D*); here, *I* denotes the set of Xenium cell that maps to a Visium spot (in the intersecting area of Visium and Xenium slices), *A*_*XI*_ and Γ_*I*_ denote *A*_*X*_ and Γ subseted to *I* respectively, *V*_*g*_ denotes the Visium expression for the holdout gene, and 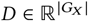 is the combination coefficients for imputing the holdout gene. We use the alignment as described in Section 3.1, and identical mapping Γ as used in SIID. We impute the expression for all Xenium cells (inside or outside *I*) after training and evaluate correlation on all cells.
- For Baseline A, the estimated expression for a Xenium gene *g* is given by Γ*V*_*g*_, where *V*_*g*_ refer to the expression of gene *g* in Visium. Intuitively, the Visium spot expression is copied to each Xenium cell mapped to that spot through Γ. Xenium cells that are not mapped to a Visium spot (as the datasets are partially aligned) are assigned zero expression.
- For Baseline B, let 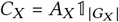 be the observed total count per Xenium cell. If a Xenium cell *i* is mapped to Visium spot *j*, we let 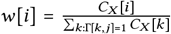 be the weight of cell *i* among all cells mapped to the same spot. The estimate for a Xenium gene *g* is then given by computing the estimate for Baseline A, then scale by *w*[*i*] for each cell *i*. Intuitively, this means the counts in Visium spot *j* is distributed among Xenium cells mapped to that spot proportional to observed Xenium counts.
- For Baseline C, following Baseline A where we copy expression of each Visium spot to all Xenium cells it maps to, we apply a kNN smoothing where the expression of each Xenium cell is calculated as the average of its 120 nearest neighbors. We then follow the partitioning step in Baseline B, scaling the expression of each cell further by its observed total Xenium count over all observed Xenium mapped to the same Visium spot.
- Baseline D is implemented in a similar fashion as Baseline C, but instead of applying the kNN smoothing before rescaling, we apply it directly on Baseline B results.

#### C.5 Evaluating the effect of noise in spatial mapping

We follow the identical setup as in Appendix B.3 to evaluate the effect of changing Γ in imputation performance. See Figure S2C and Figure S2D for their effect on imputing real datasets.

#### C.6 Consolidated runtime information for imputation runs

SIID are benchmarked on a single Tesla P100 GPU with 12GB memory, and most other methods (if CPU-Bound) are run on a cluster with 20 Xeon E5-2690 CPUs. SIID is run five times per fold with 3 restarts each and the reported values in Table 1 and Table S2 are averaged across 5 runs. Average runtime for the benchmarked methods for each fold in seconds can be found in Table S2, including Visium pairing and scRNA-seq pairing. Without the 2× downsamping in CRC scRNA-seq dataset, Tangram cannot finish within 2 days or 172,800 seconds. For Tangram in cluster mode, we do not include the time taken for running Leiden clusters.

**Table S2:** Comparison of average run time in seconds per fold for the benchmarked methods.

| Dataset | SIID (3 runs) | Tangram (Visium) |  | SLAT | SANTO | Tangram (scRNA-seq) |  |
| --- | --- | --- | --- | --- | --- | --- | --- |
|  |  | cell mode | cluster mode |  |  | cell mode | cluster mode |
| BRCA | 403s | 8,632s | < 300s | < 300s | 4,020s | 43,309s | < 300s |
| CRC | 965s | 17,232s | < 300s | 429s | 12,814s | 65,240s | < 300s |

We note that Tangram in cell mode takes very long time to complete, in part because we do not have enough GPU memory to place the Tangram model there: Tangram has at least |*S*_*X*_ ||*S*_*V*_ | parameters, while the number of parameters in SIID is around *h*(|*S*_*X*_ |+|*G*_*V*_ |) ≈ *h*|*S*_*X*_ | given |*S*_*X*_ |≫ |*G*_*V*_ |, magnitudes lower than what Tangram requires (around 100× lower for our benchmarking datasets).

#### C.7 SIID with Negative Binomial Counts

##### Modeling

Inspired by gimVI [27], we implement an alternative model where we assume that Xenium counts still follow the same Poisson distribution, while Visium counts follow a negative binomial distribution. The negative binomial mean 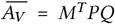 is the same as in the count-scaled reparameterization described in Section 2.2. The overdispersion *α*_*V*_ is gene-specific but consistent between spots. Formally, the loss function is

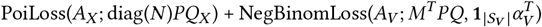

Here, 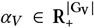 is the overdispersion for each gene, and the matrix 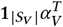 indicates that all spots for the same gene share the same overdispersion parameter. For a given spot-gene pair, with observed count *X* and inferred mean *m* and overdispersion *α*, the loss function (negative log-likelihood) is defined as:

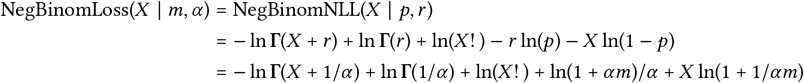

where *r* = 1*/α* and *p* = 1*/*(1 + *αm*) are the parameters of the negative binomial in its natural parameterization, and Γ is the Gamma function that extends factorial to complex numbers.

##### Implementation

We implement this version of SIID, optimizing *α*_*V*_ alongside other parameters such as *P, Q* and *M*. Since overdispersion cannot be negative, we similarly perform gradient descent on log *α*_*V*_ instead.

##### Evaluation

We evaluate this version of SIID on two real datasets in the identical setup as described in Section 3.3. It achieves a similar performance on BRCA dataset (average gene-wise *R*^2^ of 0.2498, compared to 0.2527 of original version) and slightly worse on CRC dataset (average gene-wise *R*^2^ of 0.2013, compared to 0.2248 of original version). Evaluation on other metrics can be found in Table S4 under the column “NegBinom”.

### D. Supplemental Results

#### D.1 Reporting other correlation measures

In addition to *R*^2^ scores, we use four other correlation scores as described in [23]. As shown in Table S3, SIID has the best performance by all listed measures.

With results from Tangram paired with scRNA-seq as well as SIID with Negative Binomial counts, SIID with Poisson counts model has the best performance in all but one measure where it’s second best.

**Table S3:** Comparing correlation scores on selected methods for imputing Xenium gene expression from paired Visium data. Arrow following metric name indicates whether higher is better (↑) or lower is better (↓). Bold indicates best performance in each metric and dataset pair.

| Metric | Dataset | SIID | Tangram |  | SLAT | SANTO |
| --- | --- | --- | --- | --- | --- | --- |
|  |  |  | cell mode | cluster mode |  |  |
| $R^2 \uparrow$ | BRCA | **0.2527** | 0.1874 | 0.1557 | 0.0688 | 0.1042 |
|  | CRC | **0.2248** | 0.1789 | 0.1382 | 0.0608 | 0.0727 |
| PCC ↑ | BRCA | **0.4470** | 0.3789 | 0.3402 | 0.2015 | 0.2620 |
|  | CRC | **0.4099** | 0.3644 | 0.2995 | 0.1997 | 0.2236 |
| SSIM ↑ | BRCA | **0.3456** | 0.3112 | 0.1904 | 0.1192 | 0.2239 |
|  | CRC | **0.4191** | 0.3905 | 0.2761 | 0.1017 | 0.3437 |
| RMSE ↓ | BRCA | **1.0261** | 1.0952 | 1.1317 | 1.2542 | 1.2027 |
|  | CRC | **1.0609** | 1.1089 | 1.1658 | 1.2596 | 1.2399 |
| JS ↓ | BRCA | **7.1959** | 8.3165 | 8.3599 | 12.4704 | 8.9354 |
|  | CRC | **8.2665** | 9.1641 | 9.2956 | 12.4493 | 9.8258 |

#### D.2 Supplemental Figures

**Table S4:** Similar to Table S3, with scRNA-seq pairing results from Tangram and SIID with Negative Binomial counts (Appendix C.7) also reported.

| Metric | Dataset | SIID |  | Tangram (Visium) |  | SLAT | SANTO | Tangram (scRNA-seq) |  |
| --- | --- | --- | --- | --- | --- | --- | --- | --- | --- |
|  |  | Poisson | NegBinom | cell | cluster |  |  | cell | cluster |
| $R^2 \uparrow$ | BRCA | **0.2527** | 0.2498 | 0.1874 | 0.1557 | 0.0688 | 0.1042 | 0.2194 | 0.2203 |
|  | CRC | **0.2248** | 0.2013 | 0.1789 | 0.1382 | 0.0608 | 0.0727 | 0.1960 | 0.1823 |
| PCC ↑ | BRCA | **0.4470** | 0.4431 | 0.3789 | 0.3402 | 0.2015 | 0.2620 | 0.4126 | 0.4176 |
|  | CRC | **0.4099** | 0.3786 | 0.3644 | 0.2995 | 0.1997 | 0.2236 | 0.3794 | 0.3700 |
| SSIM ↑ | BRCA | 0.3456 | 0.3429 | 0.3112 | 0.1904 | 0.1192 | 0.2239 | **0.4930** | 0.2994 |
|  | CRC | 0.4191 | 0.3953 | 0.3905 | 0.2761 | 0.1017 | 0.3437 | **0.5786** | 0.3494 |
| RMSE ↓ | BRCA | **1.0261** | 1.0296 | 1.0952 | 1.1317 | 1.2542 | 1.2027 | 1.0609 | 1.0575 |
|  | CRC | **1.0609** | 1.0900 | 1.1089 | 1.1658 | 1.2596 | 1.2399 | 1.0926 | 1.1038 |
| JS ↓ | BRCA | **7.1959** | 7.2051 | 8.3165 | 8.3599 | 12.4704 | 8.9354 | 7.8144 | 7.8562 |
|  | CRC | **8.2665** | 8.4766 | 9.1641 | 9.2956 | 12.4493 | 9.8258 | 8.8543 | 8.8650 |

**Figure S1:**
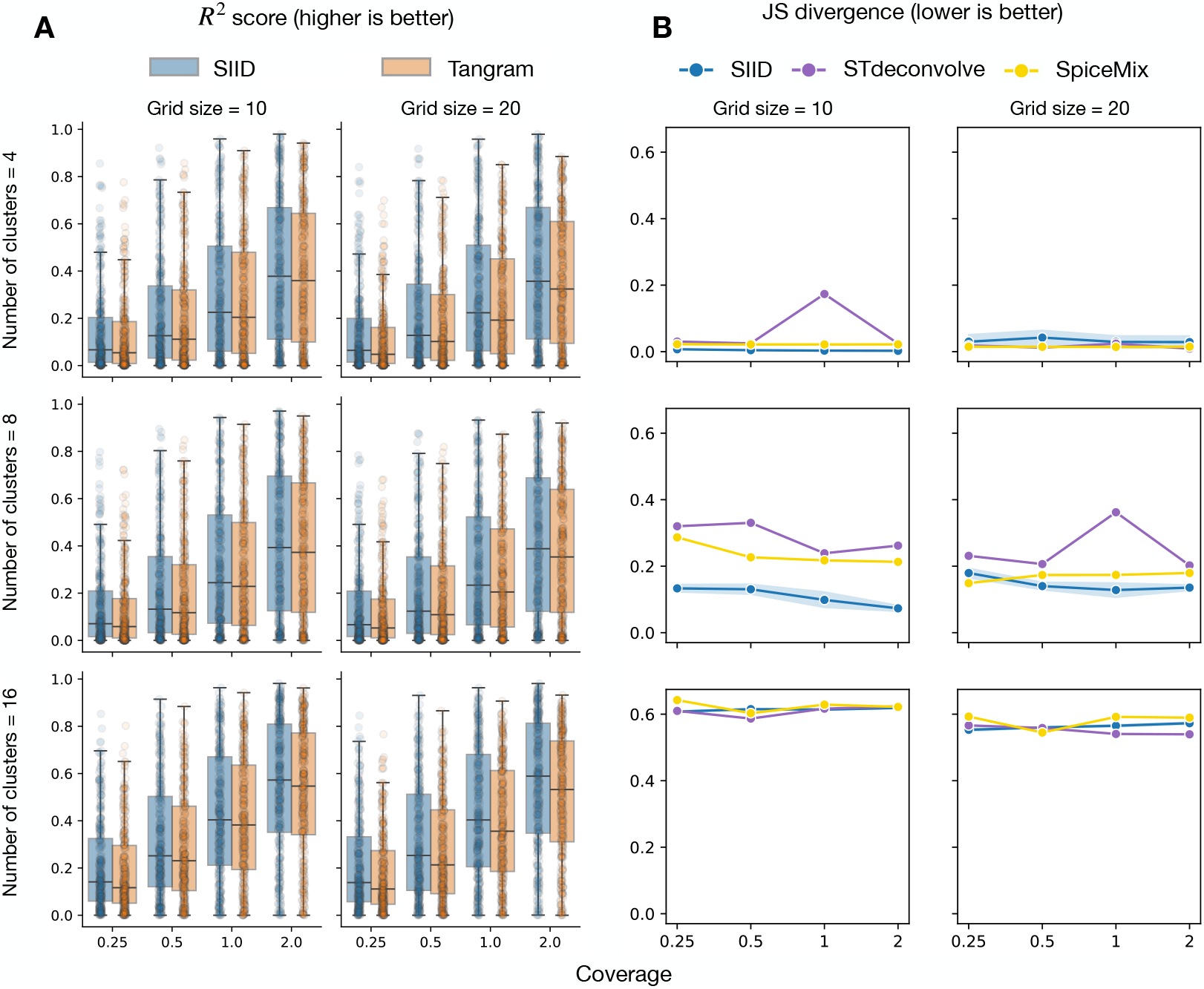
**A:** *R*^2^ score for each gene’s expression between predicted values (from SIID and Tangram) and ground truth across different simulation settings. **B:** Jensen-Shannon (JS) divergence of predicted cell type mixture proportions compared to the ground truth for SIID, STdeconvolve and SpiceMix.

**Figure S2:**
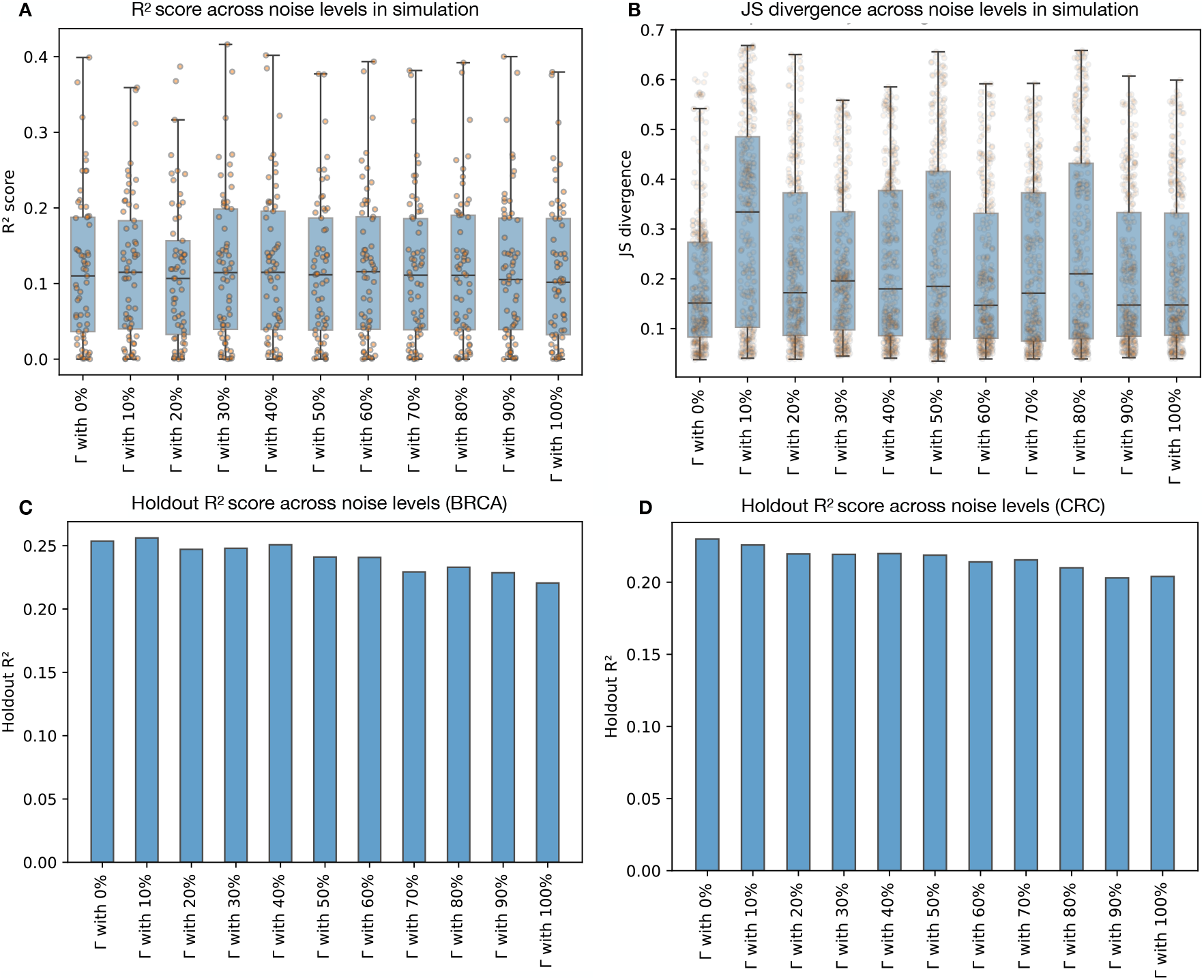
Evaluating model performance in the presence of perturbed Γ. **A:** *R*^2^ score of imputed holdout genes across different level of noise in simulated scenarios. Each point indicate a gene. **B:** JS divergence across Visium spots across different level of noise in simulated scenarios. Each point indicate a Visium spot in simulation. **C:** Average *R*^2^ score of imputed holdout genes across different level of noise in BRCA holdout experiments. **D:** Same as panel C over the CRC dataset.

**Figure S3:**
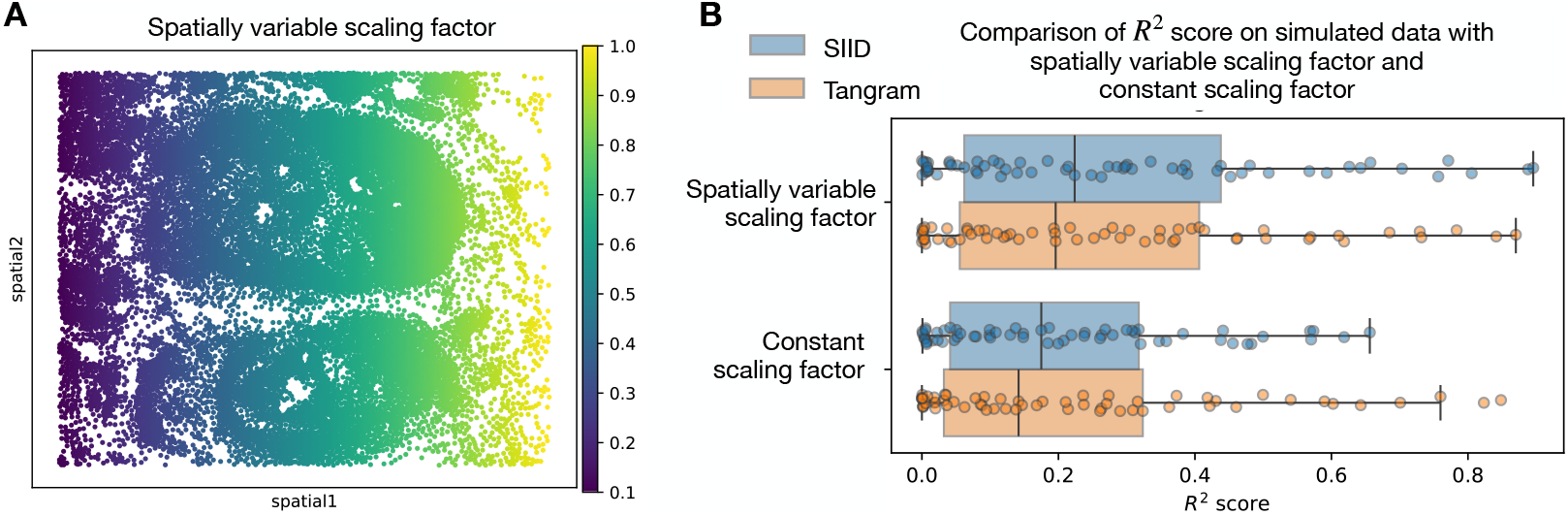
Evaluating the effect of spatially variable scaling factor in simulation. **A:** Visualization of spatially variable scaling factor as a multiplier for UMI count per simulated Xenium cell. **B:** Comparing *R*^2^ scores of SIID with spatially variable scaling factor and without it, with similar amount of total UMIs in each simulation.

**Figure S4:**
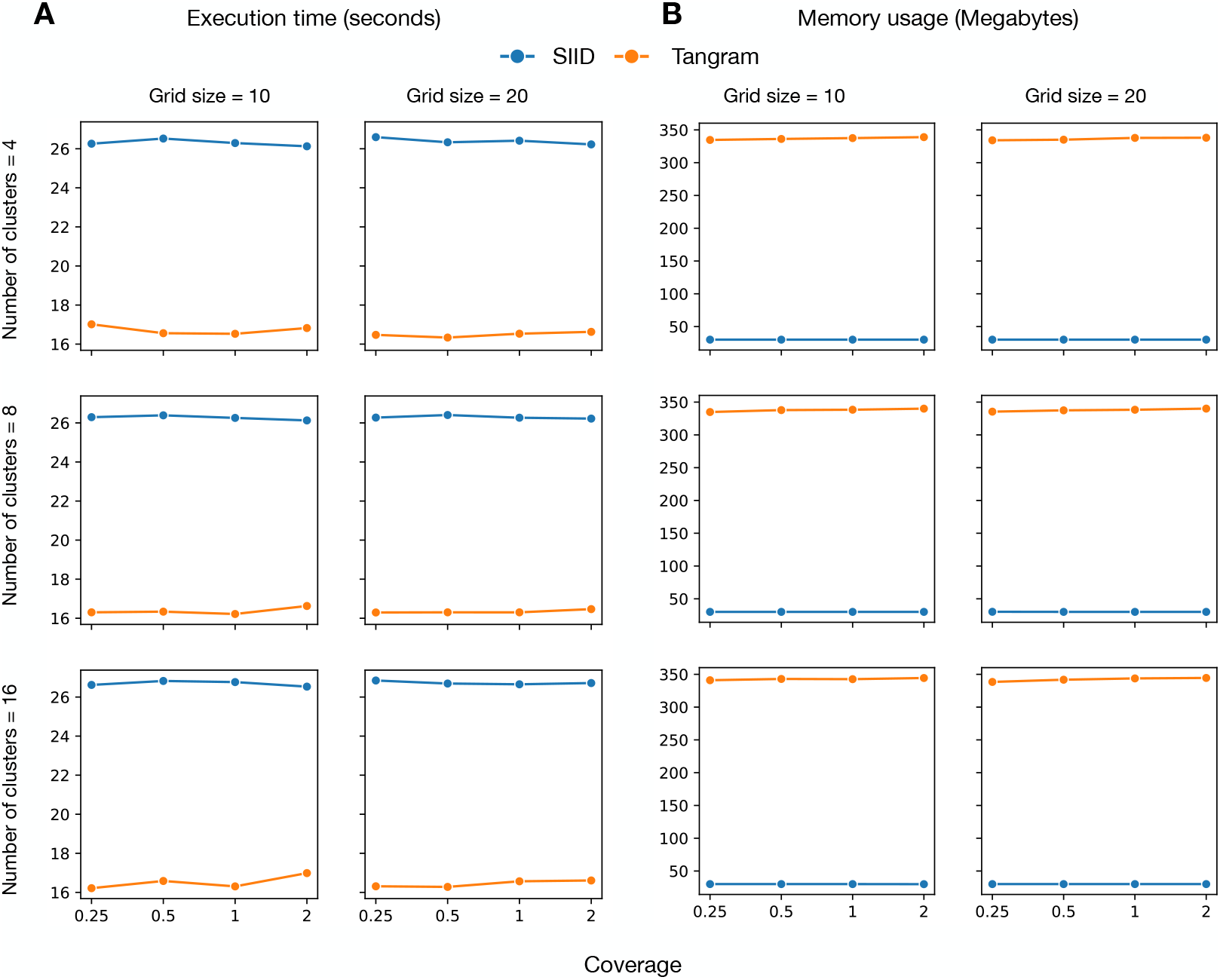
Computational performance comparison of SIID and Tangram across different simulation configurations. **A:** Execution time (in seconds) for SIID (blue) and Tangram (orange) across varying number of clusters (4,8,6), grid sizes (10, 20), and coverage levels (0.25, 0.5, 1, 2). **B:** Memory usage in megabytes for SIID (blue) and Tangram (orange) under the same simulation setting.

**Figure S5:**
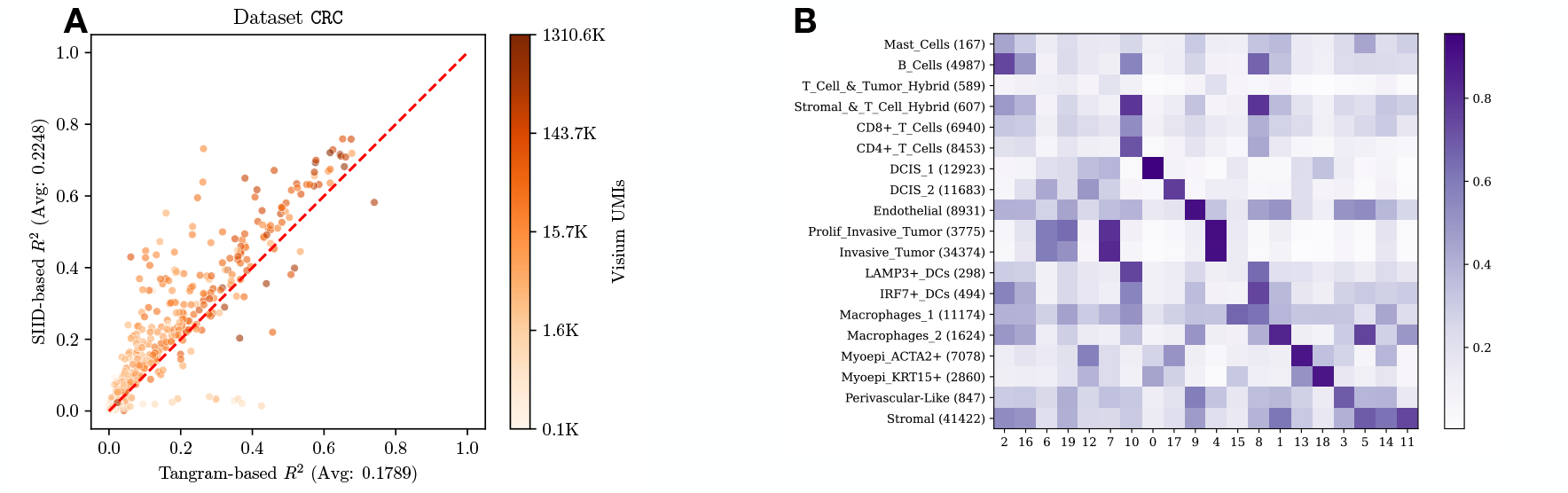
**A:** Per-gene *R*^2^ score plot for CRC dataset. **B:** For Visium deconvolution, cosine similarity between inferred cluster proportions from our method (columns) and RCTD inferred cell type proportions (rows). Note that unlabeled cells are discarded when running RCTD.

**Figure S6:**
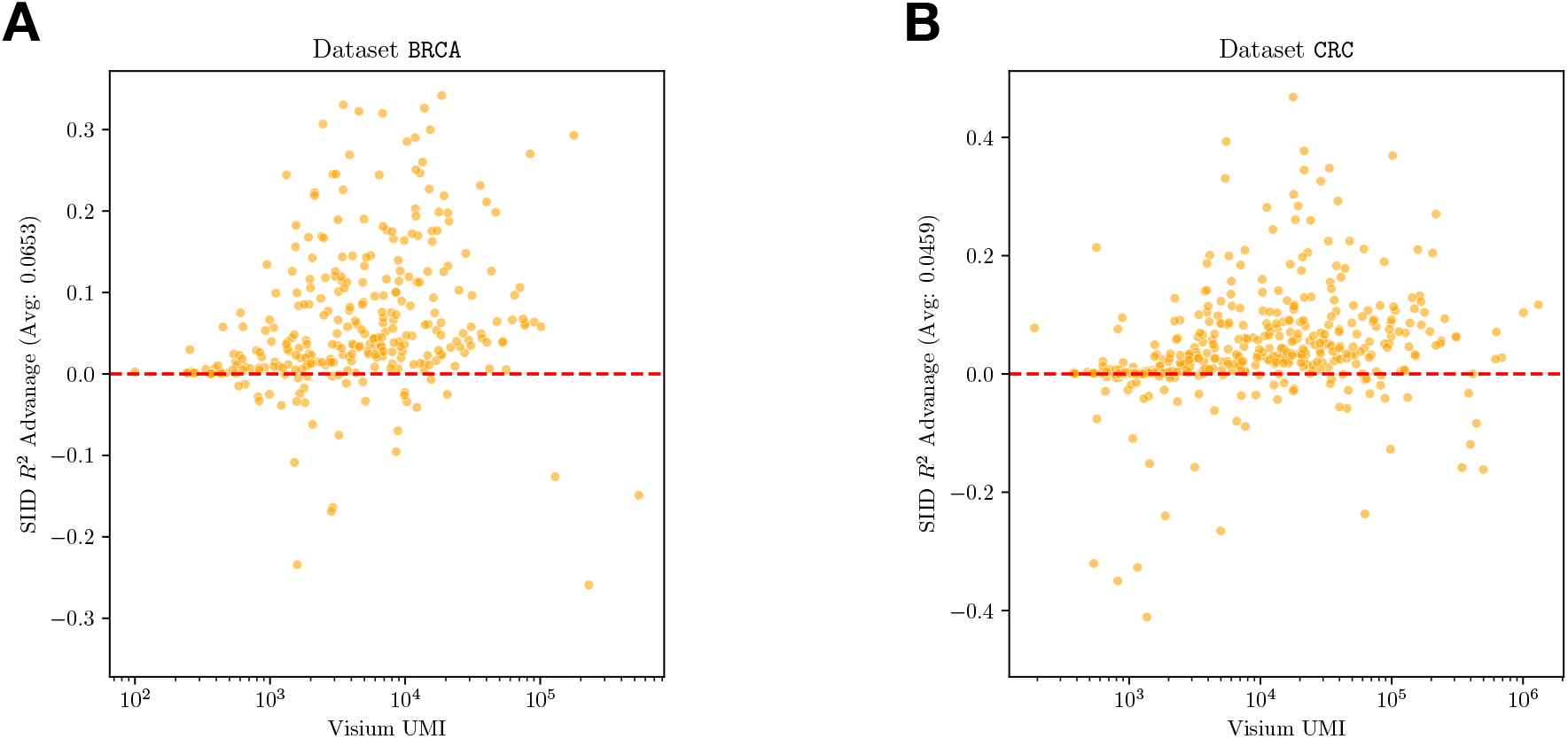
*R*^2^ score differential for each individual gene in both **A:** BRCA dataset and **B:** CRC dataset. X-axis represents log-scaled total UMI counts for each gene in Visium. Red dotted line represents *y* = 0, dots above the line denotes SIID achieving higher *R*^2^ compared to Tangram for that gene.

**Figure S7:**
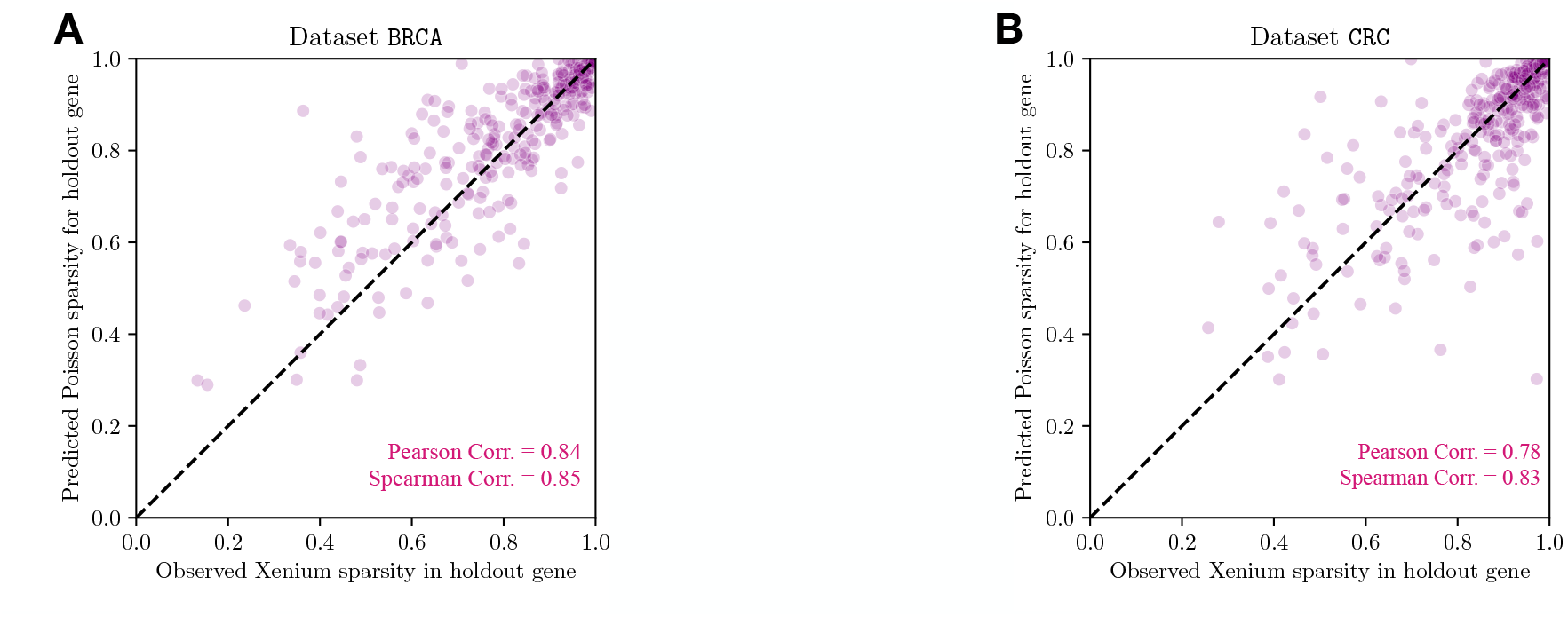
SIID recovers sparsity of Xenium gene expression in **A:** BRCA dataset and **B:** CRC dataset. X-axis represents observed sparsity of each gene in the Xenium dataset. Y-axis represents predicted sparsity of each gene in the holdout experiment given *P* (*Y* = 0 | *Y* ~ Pois(*X*)) = *e* ^−*X*^ for each observation. Bottom right indicates Pearson and Spearman correlation of sparsity prediction.

**Figure S8:**
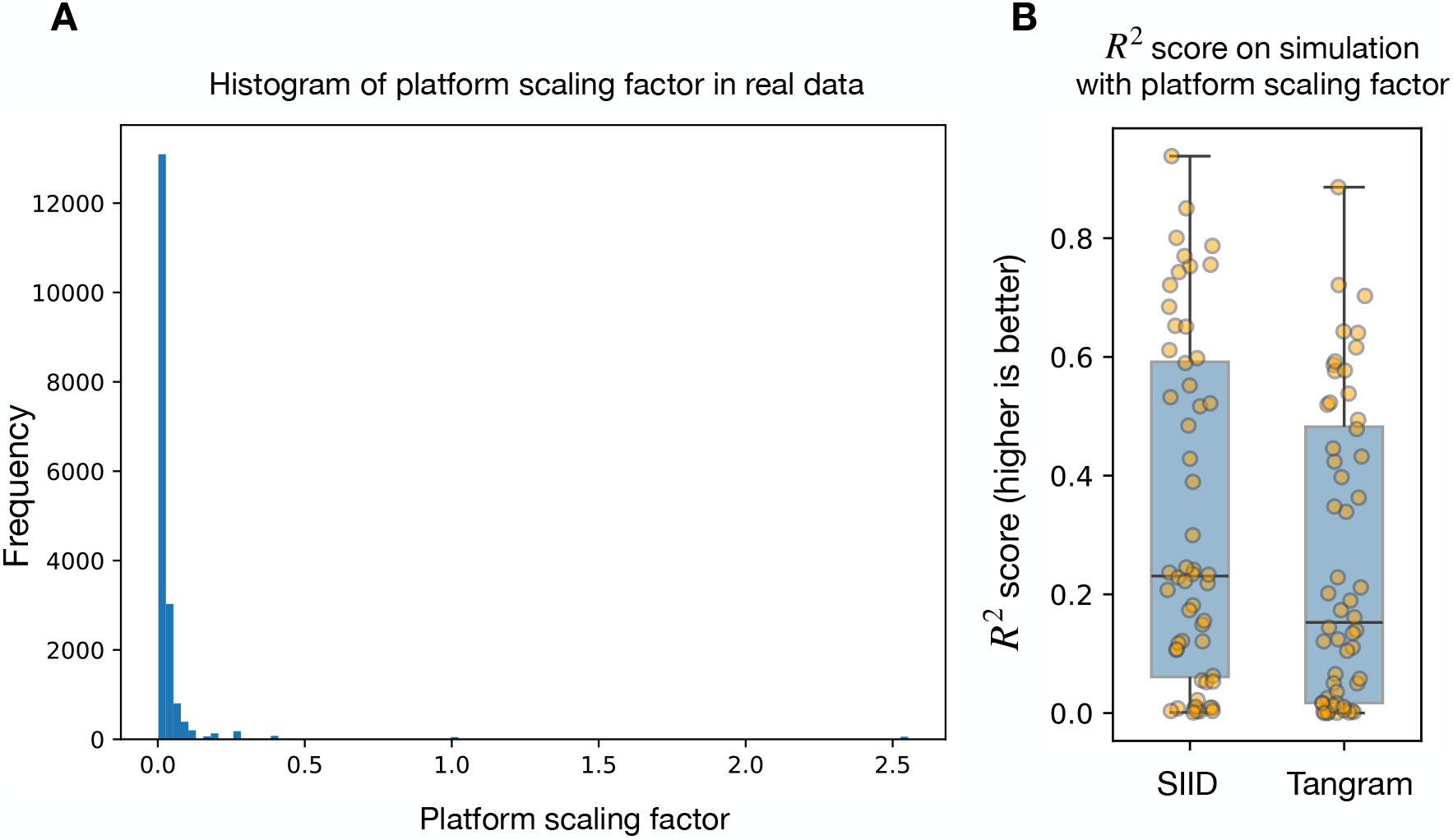
**A:** Histogram of platform specific scaling factor for shared genes from BRCA. **B:** The *R*^2^ score for imputed expression on holdout genes in simulated data with platform specific scaling from SIID and Tangram.

**Figure S9:**
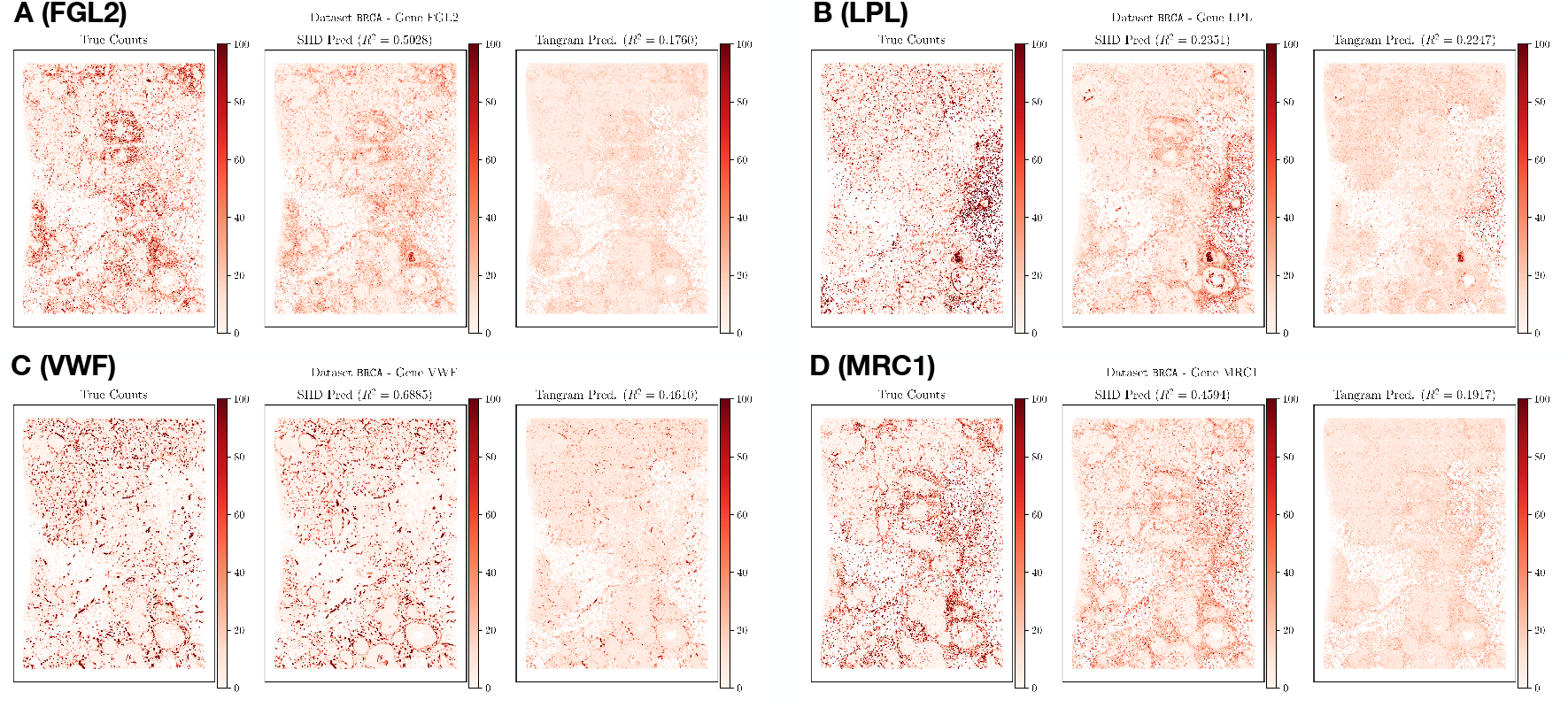
Left: Observed Xenium expression (ground truth for evaluating imputation) Middle: SIID prediction when the gene is held out, Right: Tangram prediction for **A** FGL2, **B** LPL, **C** VWF, and **D** MRC1 genes when they are held out. We also present the *R*^2^ scores between predicted expression and observed expression in the title. We normalize the total counts within each gene to 1, 000, 000 and plot all normalized counts on the same color scale.

## Footnotes

1 https://pytorch.org/

2 SLAT and SANTO are only applicable to SRT datasets and therefore are not included in this comparison. gimVI failed to finish on several runs (details in Appendix C.4).

3 https://github.com/JEFworks-Lab/STdeconvolve

4 https://github.com/ma-compbio/spicemix

5 https://www.10xgenomics.com/products/xenium-in-situ/preview-dataset-human-breast

6 https://www.10xgenomics.com/products/visium-hd-spatial-gene-expression/dataset-human-crc

